# Structural Mapping of the EIN2–EIN3 Interaction Core and Its Integration with ENAP1 in Ethylene Signaling

**DOI:** 10.64898/2026.07.10.737652

**Authors:** Fabian Wynen, Meik Thiele, Maya Hettesheimer, Raphael Josef Eberle, Jan Eric Maika, Rüdiger Simon, Georg Groth

## Abstract

Ethylene regulates diverse developmental processes, yet the molecular function of its central regulator, ETHYLENE INSENSITIVE 2 (EIN2), has remained unclear. Although EIN2 nuclear import is mediated by the Importin-α/β pathway, the molecular events initiated by EIN2 after nuclear entry were unknown. Here we show that EIN2 directly engages the transcription factor EIN3, establishing a mechanistic link between EIN2 nuclear accumulation and transcriptional activity. Microscale thermophoresis, yeast two-hybrid analysis and *in planta* FLIM-FRET consistently support this interaction. Domain mapping identifies EIN3 residues 86-173 as the core EIN2-binding region, and structural modeling refines the interface to a conserved segment within residues 86-120 that contacts a conserved region near the N-terminus of the EIN2-CEND fragment. *In planta*, EIN2 residues 1042-1214 are sufficient for EIN3 binding, revealing multiple interaction-competent surfaces with distinct affinities. The chromatin-associated protein ENAP1 also binds EIN2 and competes with EIN3, indicating a dynamic, concentration-dependent regulatory mechanism rather than a stable ternary complex. These findings define the molecular basis of the EIN2-EIN3 interaction and provide a mechanistic framework for EIN2-dependent transcriptional control in ethylene signaling.

## INTRODUCTION

Ethylene is a gaseous plant hormone that regulates a broad spectrum of developmental, and stress-related processes, including inhibition of cell elongation, seed germination, apical hook formation, senescence, fruit ripening, abscission, and responses to biotic and abiotic stresses^1,2^. Ethylene perception and signaling occur through a largely linear pathway that transmits information from the endoplasmic reticulum (ER) membrane to the nucleus. In *Arabidopsis* five ER-localized ethylene receptors (ETR1, ETR2, ERS1, ERS2 and EIN4) act as negative regulators of the pathway^3,4^. In the absence of ethylene, these receptors activate the Raf-like kinase CONSTITUTIVE TRIPLE RESPONSE1 (CTR1), which suppresses downstream signaling. Ethylene binding inactivates receptor-CTR1 signaling, thereby relieving repression of the central regulator ETHYLENE INSENSITIVE2 (EIN2)^5,6^. EIN2 is the pivotal conduit between ER-localized ethylene perception and nuclear transcriptional responses. It is an ER-anchored protein comprising an N-terminal NRAMP-like (Natural Resistance-Associated Macrophage Proteins) domain and a large C-terminal cytosolic region. Upon ethylene perception, CTR1 inactivation prevents EIN2 phosphorylation, enabling proteolytic cleavage and nuclear translocation of the EIN2 C-terminal fragment (EIN2-C)^7–9^. EIN2-C regulates ethylene responses through both indirect and direct mechanisms. Indirectly, EIN2-C stabilizes the transcription factors ETHYLENE INSENSITIVE3 (EIN3) and EIN3-LIKE1 (EIL1) by inhibiting their ubiquitin-mediated degradation via the F-box proteins EBF1 and EBF2^10,11^. This mechanism includes the formation of processing bodies (P-bodies), where EIN2-C interacts with EIN5, nonsense-mediated decay proteins (UPFs) and the EBF1/2 mRNAs to suppress of EBF1/2 translation^12,13^.

In addition to these cytosolic functions, nuclear-localized EIN2-C interacts with EIN2 NUCLEAR-ASSOCIATED PROTEIN1 (ENAP1), a histone-binding protein implicated in chromatin remodeling. This interaction has been proposed to promote association with ethylene-responsive loci and to modulate H3K14 and H3K23 acetylation, thereby facilitating EIN3-dependent transcriptional regulation^14,15^. Among the six members of the EIN3 family (EIN3 and EIL1-5) in *A. thaliana*, EIN3 and EIL1 constitute the key regulatory hub of ethylene signaling as *ein-3 eil-1* double mutants abolish ethylene responses^16^. EIN3/EIL proteins share a conserved N-terminal DNA-binding domain (DBD) that recognizes EIN3-binding sites (EBS) in target promoters^17^. EIN3 functions as a homodimer, a state detectable even in the absence of DNA ^17,18^. The N-terminal region contains several critical structural components, including an acidic domain (AD), a proline-rich region (PR), and five clustered basic domains (BD I-V)^19,20^, whereas the C-terminal region is more divergent among species and may contribute to functional diversification.

Although previous studies proposed a model in which ethylene responses may involve a ternary EIN2-EIN3-ENAP1 complex^15^, the molecular basis of these interactions and the existence of such complex remain unresolved. Moreover, the EIN2 nuclear localization signal (NLS) has been reported to interact with ETR1 and Importin-α/β family members^9,21,22^, raising the question of whether the NLS contributes additional regulatory functions beyond nuclear import.

Here, we demonstrate direct EIN2-EIN3 binding across multiple experimental contexts and define the interface that mediates this interaction. We further refine the structural organization of EIN3 by delineating its homodimerization domain, nuclear localization requirements and functional regions. Competitive binding assays with ENAP1 reveal that EIN2 engages multiple nuclear partners, pointing to a more complex interaction landscape than previously recognized and providing new mechanistic insight into the nuclear events following EIN2 activation.

## RESULTS

### Protein-protein interaction (PPI) network analysis of EIN2

EIN2 is a central regulator of ethylene signaling and integrates multiple layers of control at both the protein and transcriptional levels. Previous studies have shown that EIN2 directly interacts with ethylene receptor ETR1, positioning it as a key mediator that transmits signals from receptor complexes to downstream components^5,21^. In addition, EIN2 promotes the formation of cytoplasmic processing bodies (P-bodies), linking it to post-transcriptional regulation through effects on mRNA stability^12^. EIN2 is further functionally connected to the transcription factor EIN3, thereby bridging upstream ethylene perception with nuclear transcription responses^7,12^ (Figure 1A).

**Figure 1:**
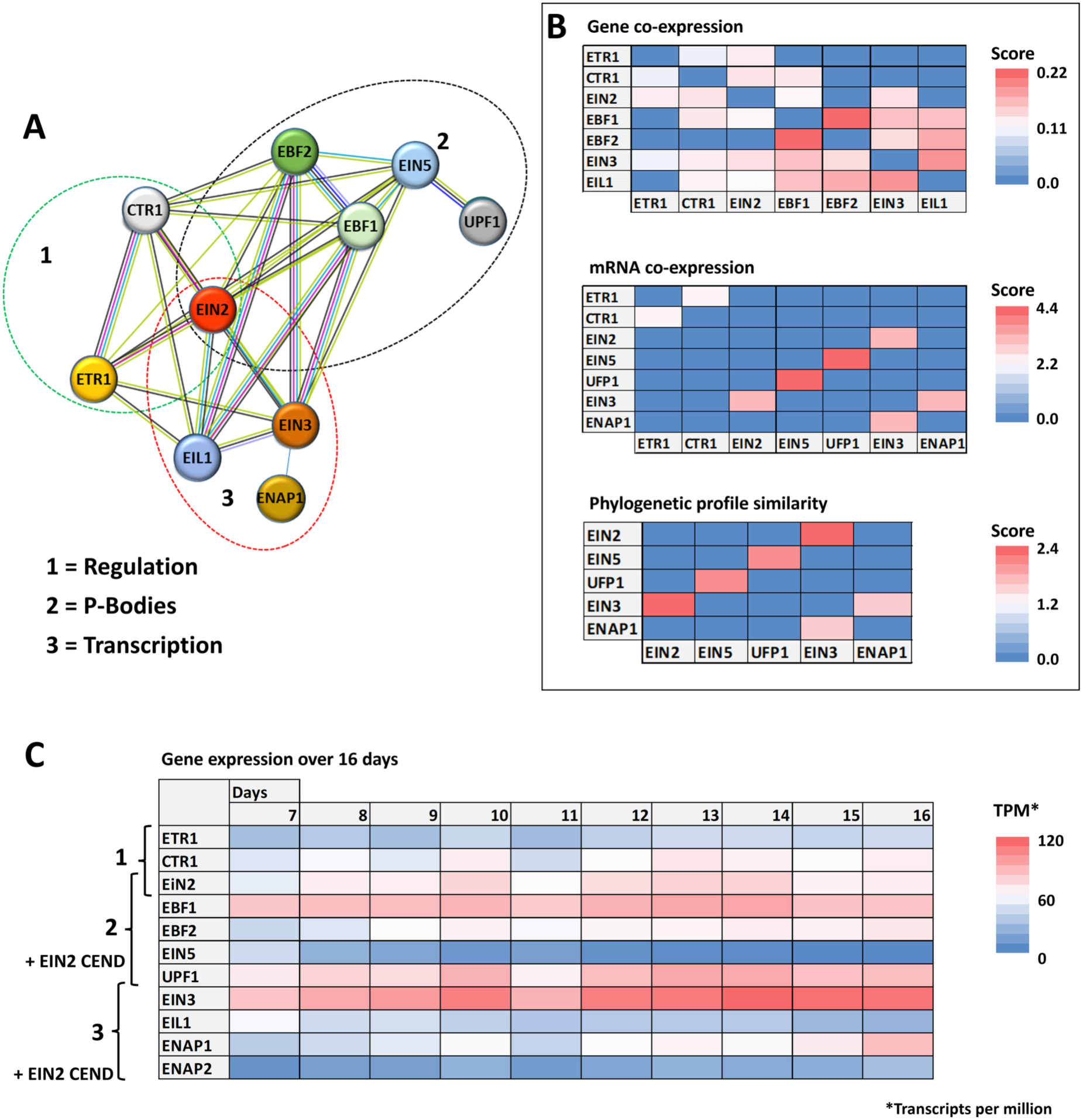
Bioinformatical analysis of ethylene-related proteins in *A. thaliana* including protein-protein interaction networks, co-expression patterns, and temporal gene expression. Analyses were performed using data from STRING^34^, FunCoup6^24^ and Expression Atlas^23^, restricted to *A. thaliana*. **(A)** Protein-protein-interaction (PPI) network, highlighting the central positions of EIN2 and EIN3 within the ethylene signaling pathway. Proteins associated with (1) ETR1-mediated receptor regulation, (2) P-body formation, and (3) transcriptional regulation are grouped to illustrate functional modules. **(B)** Heat map summarizing (top) gene co-expression of core ethylene signaling components (ETR1, CTR1, EIN2, EBF1, EBF2, EIN3 and EIL1), (middle) mRNA co-expression of proteins involved in post-transcriptional and translational regulation (ETR1, CTR1, EIN2, EIN5, UPF1, EIN3 and ENAP1), and (bottom) phylogenetic profile similarity of EIN2, EIN5, UFP1, EIN3 and EIN2, reflecting functional relationships across plant species. **(C)** Temporal expression heat map showing transcript abundance of ethylene-related genes (ETR1, CTR1, EIN2, EBF1, EBF2, EIN5, UPF1, EIN3, EIL1, ENAP1 and ENAP2) across a 16-day developmental time course in *A. thaliana*.

Co-expression analyses support this functional relationship. EIN2 and EIN3 transcripts show tightly coordinated expression patterns (Figure 1B, Expression atlas^23^), consistent with their sequential roles in ethylene signal transduction^5,20^. Quantitative expression data from FunCoup6^24^ indicate that EIN2 and EIN3 are expressed at comparable levels, suggesting balanced regulation within the pathway (Figure 1B). ENAP1 (EIN2 NUCLEAR ASSOCIATED PROTEIN 1) also displays co-expression with EIN3, in line with its proposed role as chromatin-associated co-regulator of ethylene-responsive transcription. Phylogenetic profile similarity analyses further reveal that EIN2, EIN3, and ENAP1 are deeply conserved across plant species (FunCoup6^24^). Such conservation typically reflects essential biological functions and evolutionary constraints. The strong similarity in their phylogenetic distribution suggests that these proteins may have co-evolved as components of a conserved regulatory module in ethylene signaling.

Temporal expression profiling across a 16-day developmental series in *Arabidopsis thaliana* (Expression Atlas^23^) show that EIN2 maintains relatively constant transcript levels, whereas EIN3 expression increases markedly after day 12, indicating potential age-dependent regulation. ENAP1 expression remains stable throughout the time course, consistent with a sustained role in transcriptional regulation, while ENAP2 expression remains low (Figure 1C).

Previous ChIP-seq studies demonstrated that acetylation of histones H3K23 and H3K14 increases upon ethylene treatment in an EIN2-dependent manner. EIN2 interacts with ENAP1, and ENAP1 can bind EIN3, leading to the hypothesis that ENAP1 and EIN2 jointly stabilize EIN3 and promote chromatin acetylation at ethylene-responsive loci^14,15^. Despite these insights, the molecular architecture of this regulatory module remains unresolved. In particular, it is unclear whether EIN2 directly interacts with EIN3 or whether ENAP1 serves as a bridging factor to form a ternary complex.

To address these open questions, we employed a combined set of biophysical and cell-based assays across multiple biological contexts to determine whether EIN2 directly interacts with EIN3 and to dissect the interaction interfaces among EIN2, EIN3 and ENAP1. These analyses provide critical mechanistic insight into the protein interaction network that governs nuclear ethylene signaling.

### Microscale thermophoresis (MST)-based characterization of EIN2–EIN3 and EIN2/EIN3-ENAP1 interactions

To dissect the molecular interfaces underlying the nuclear interactions of EIN2, we recombinantly expressed and purified a series of EIN2 C-terminal truncations (residues 479-1294, 663-1294, and 1078-1294), EIN3 N-terminal truncations (residues 1-85, 1-173, 1-306 and 1-471), and full-length ENAP1 (Figure 2A, Figure 3A). Microscale thermophoresis (MST) measurements revealed that EIN2 directly binds EIN3 with high affinity (K_D_ = 0.27 - 4.8 µM; Figure 2B).

**Figure 2:**
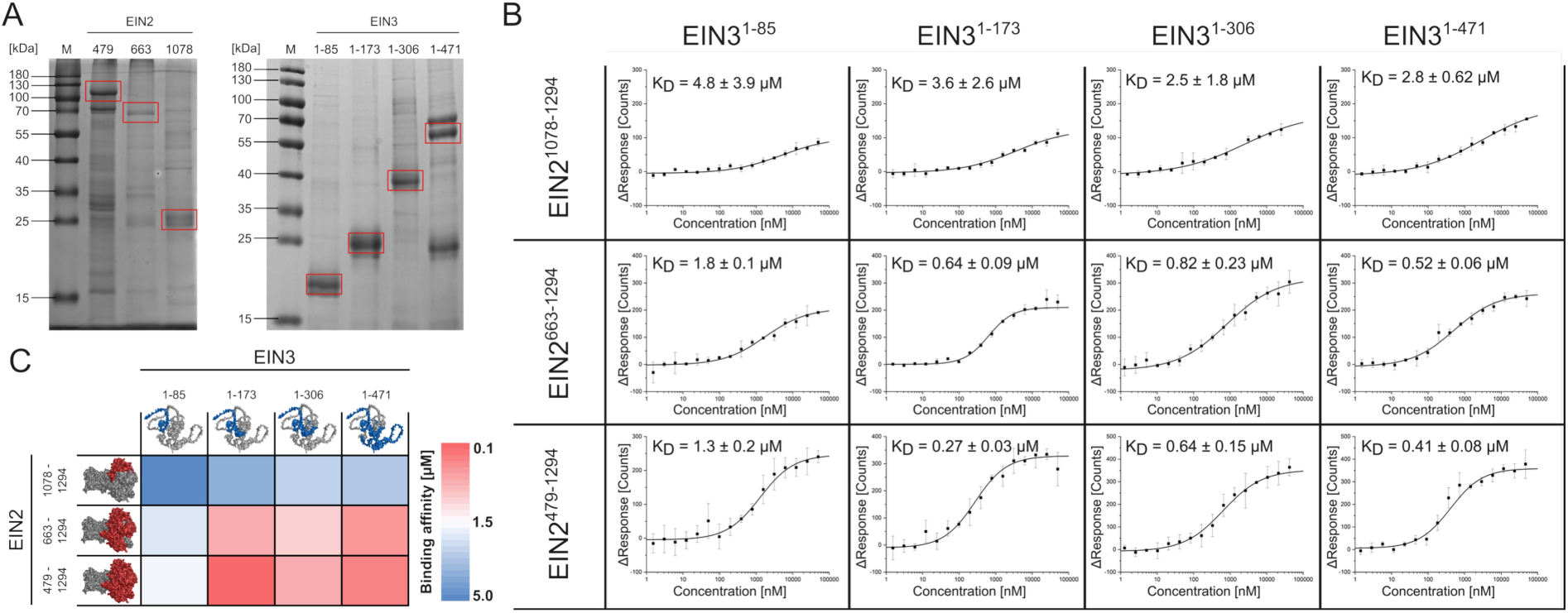
*In vitro* MST analysis of EIN2 and EIN3 truncation binding. **A)** SDS-PAGE and Coomassie staining of purified EIN2 and EIN3 truncations. The corresponding protein bands for each construct are indicated by red boxes. **B)** Microscale Thermophoresis (MST) binding curves and dissociation constants (K_D_, µM) for EIN2 and EIN3 truncations. EIN2 constructs were labeled with Alexa488 and used as fluorophores. Binding data were evaluated using initial fluorescence analysis, and curves were fitted with a logistic model in Origin. Data represent mean +/− standard deviation from three independent measurements (n = 3). **C)** Heat-map representation of KD values (µM) for all EIN2-EIN3 truncation combinations shown in B), with higher affinity depicted in red and lower affinity in blue. For visualization, structural models of EIN2 and EIN3 were generated using Chai Discovery^40^ or AlphaFold3^41^, respectively, and the amino acid regions corresponding to each truncation are highlighted in red (EIN2) or blue (EIN3).

**Figure 3:**
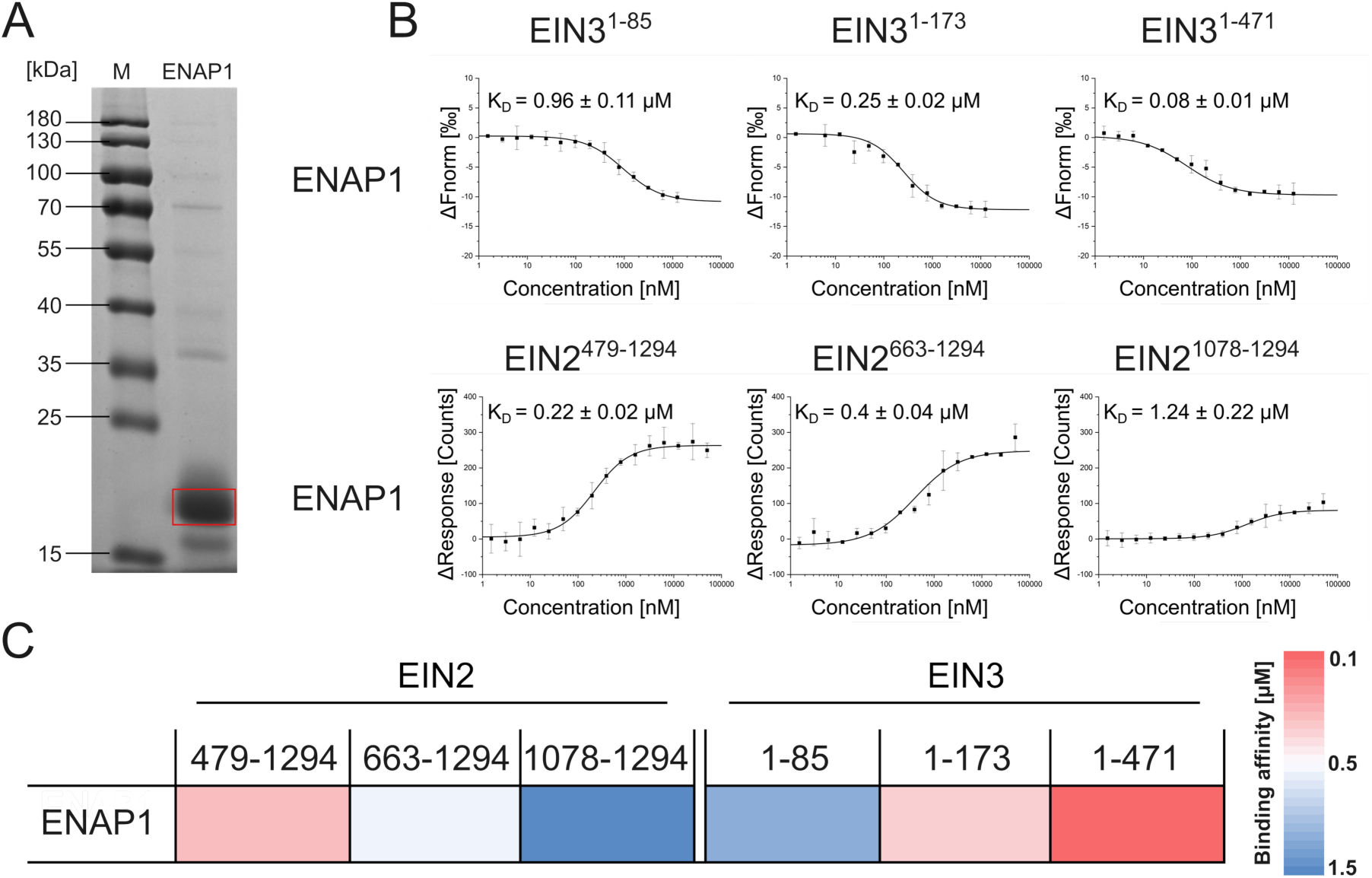
MST-based analysis of ENAP1 binding to EIN2 and EIN3 truncations. **A)** SDS-PAGE and Coomassie staining of purified ENAP1. The ENAP1 band is indicated by red boxes. **B)** Microscale Thermophoresis (MST) binding curves showing affinities (Kd, µM) between ENAP1 and EIN2 or EIN3 truncations. EIN2 truncations were labeled with Alexa488 for ENAP1 binding assays, whereas ENAP1 served as the fluorophore in EIN3 binding experiments. Binding data were evaluated using Initial Fluorescence (ENAP1-EIN2) or MST trace analysis (ENAP1-EIN3) and fitted with a logistic model in Origin. Data represent mean +/− SD from three independent measurements (n = 3). **C)** Heatmap representation of (K_D_) values (µM) for ENAP1 binding to EIN2 and EIN3 truncations. Higher affinity is shown in red, lower affinity in blue.

EIN3^1–85^ displayed only weak binding to all EIN2 truncations, whereas longer EIN3 constructs showed markedly increased affinity, particularly towards EIN2^479–1294^ and EIN2^663–1294^. EIN3^1–173^ exhibited the highest affinity and further extension of the EIN3 C-terminus did not substantially enhance binding. These data identify EIN3 residues 86-173 as the core region required for EIN2 interaction. Consistent with this, affinities decreased progressively with shorter EIN2 constructs. Negative controls using BSA or Alexa488 dye showed no detectable interaction confirming the specificity of the EIN2-EIN3 association (Figure S1).

We next examined ENAP1 binding. MST analysis confirmed that ENAP1 interacts with both EIN2 and EIN3 (Figure 3B). As observed for EIN2-EIN3 binding, ENAP1 affinity toward EIN2 decreased with shorter EIN2 constructs: a modest two-fold reduction for EIN2^663–1294^ (0.4 µM ± 0.04) and a six-fold reduction for EIN2^1078–1294^ (1.24 µM ± 0.22) relative to EIN2^479–1294^ (0.22 µM ± 0.02).

ENAP1 also bound all EIN3 truncations with low-to-mid micromolar affinity. Extension of the EIN3 86-173 region increased affinity four-fold, and further extension to residues 307-471 produced an additional three-fold increase, suggesting that ENAP1 engages multiple regions within EIN3 (Figure 3B). To test whether ENAP1 modulates EIN2-EIN3 complex formation, we performed competition assays using EIN2^479–1294^ and EIN3^1–173^, the constructs with the highest intrinsic affinity (Figure 4). At ENAP1 concentrations approximating its K_D_ (∼ 0.2 µM) EIN2-EIN3 binding increased approximately two-fold. In contrast, a 5-fold excess of ENAP1 (∼ 1 µM) markedly reduced MST response amplitude, revealing a dynamic interaction landscape among these nuclear ethylene signaling components.

**Figure 4:**
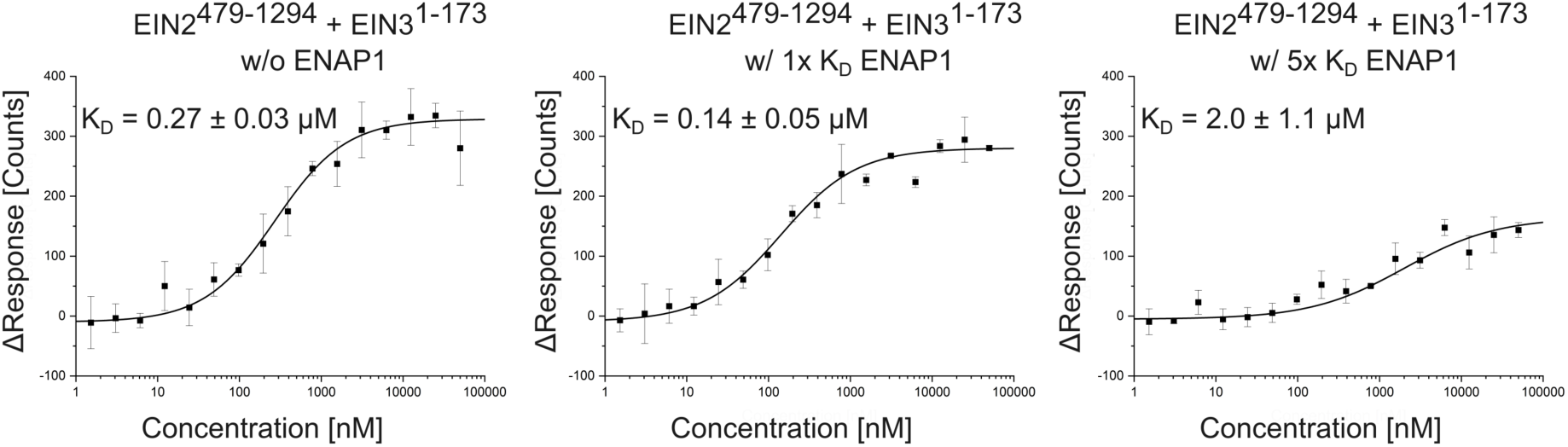
Competitive MST analysis of EIN2-EIN3 binding in the presence of ENAP1. Microscale Thermophoresis (MST) binding curves showing the interaction between EIN2^479–1294^ (Alexa488-labeled) and EIN3^1–173^ in the absence or presence of ENAP1. ENAP1 was added at concentrations corresponding to its observed binding affinity towards EIN2^479–1294^ (1x K_D_ = 0.2 µM; 5 x K_D_ = 1 µM). Binding data were evaluated using Initial Fluorescence analysis and fitted with a logistic model in Origin. Data represent mean +/− SD from three independent measurements (n = 3).

### Yeast two-hybrid (Y2H) analysis confirms the EIN2-EIN3 interaction in a cellular context

To validate the direct EIN2-EIN3 interaction observed *in vitro* in a cellular context, we next performed yeast two-hybrid (Y2H) studies using the *Saccharomyces cerevisiae* strain AH109 (Figure 5A-B). EIN2^479–1294^ was fused to the GAL4 DNA-binding domain (BD), whereas EIN3 truncations were fused to the GAL4 activation domain (AD) to avoid the known auto-activation abilities of EIN3-BD^25,26^. Successful co-transformation was confirmed on SD-LW medium, and protein-protein interactions were assessed on SD-LWH agar plates supplemented with 0.5 mM 3-AT (3-amino-1,2,4-triazole). Empty vectors served as negative controls, and the bHLH039-Fit-C interaction was used as a positive control^27^. Consistent with MST measurements, BD-EIN2^479–1294^ and full-length AD-EIN3 supported growth on SD-LWH, demonstrating a direct interaction *in vivo*. BD-EIN2^479–1294^ alone exhibited moderate auto-activation, but co-expression with AD-EIN3 resulted in visible stronger growth, confirming a genuine interaction despite the background activity. Interaction was retained with EIN3^1–173^ but not with EIN3^1–85^ supporting the conclusion that EIN3 residues 86-173 constitute the core EIN2 binding region, in agreement with MST data. Notably, EIN3^1–85^ suppressed the auto-activation of BD–EIN2^479–1294^, indicating that this short EIN3 fragment interferes with the reporter activity of the BD-fusion. This effect is unlikely to reflect productive EIN2 binding and may instead arise from non-specific interference with BD–EIN2 function in yeast, for example through partial misfolding, aggregation, or steric hindrance. Further truncation of EIN2 eliminated both auto-activation and interaction with EIN3 (Figure 5B). In particular, BD-EIN2^663–1294^ failed to interact with EIN3 despite only minor affinity differences between EIN2^663–1294^ and EIN2^479–1294^ *in vitro*, suggesting that the EIN2 region between residues 479-663 is required for productive interaction in the cellular context. The lack of EIN2^663–1294^ interaction in yeast likely reflects folding or localization constraints rather than absence of binding.

**Figure 5:**
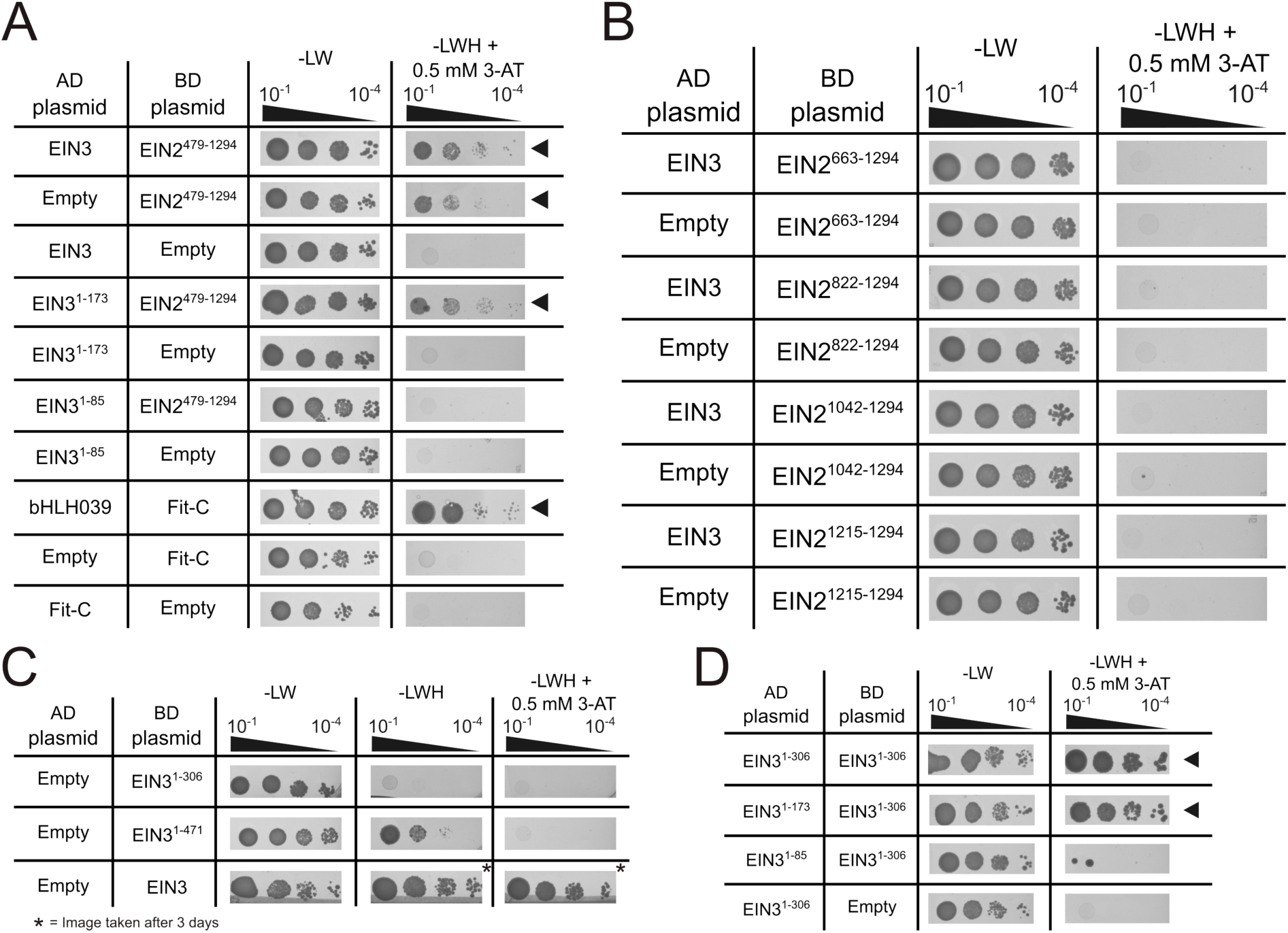
Yeast Two Hybrid (Y2H) analysis of EIN2-EIN3 interactions and EIN3 dimerization. **A)** Interaction between EIN2^479–1294^ fused to the GAL4 binding domain (BD) and EIN3 truncations fused to the GAL4 activation domain (AD). AH109 yeast co-transformants were spotted in 10-fold serial dilution (OD_600_ = 10^−1^ – 10^−4^) on SD-LW medium (transformation control) and SD-LWH medium (interaction selection) supplemented with 0.5 mM 3-AT. Images were taken after 3 days (SD-LW), or 8 days (SD-LWH). The bHLH039-Fit-C pair served as positive control as previously reported^27^, while empty vectors served as negative controls. Arrows indicate positive interactions. **B)** Interaction analysis of AD-EIN3 and BD-EIN2 truncations, performed as described in panel A. **C)** Auto-activation assays of BD-EIN3 truncations. **D)** EIN3 dimerization analysis. BD-EIN3^1–306^ was transformed with different AD-EIN3 truncations and tested as described in panel A. Arrows indicate interactions.

**Figure 6:**
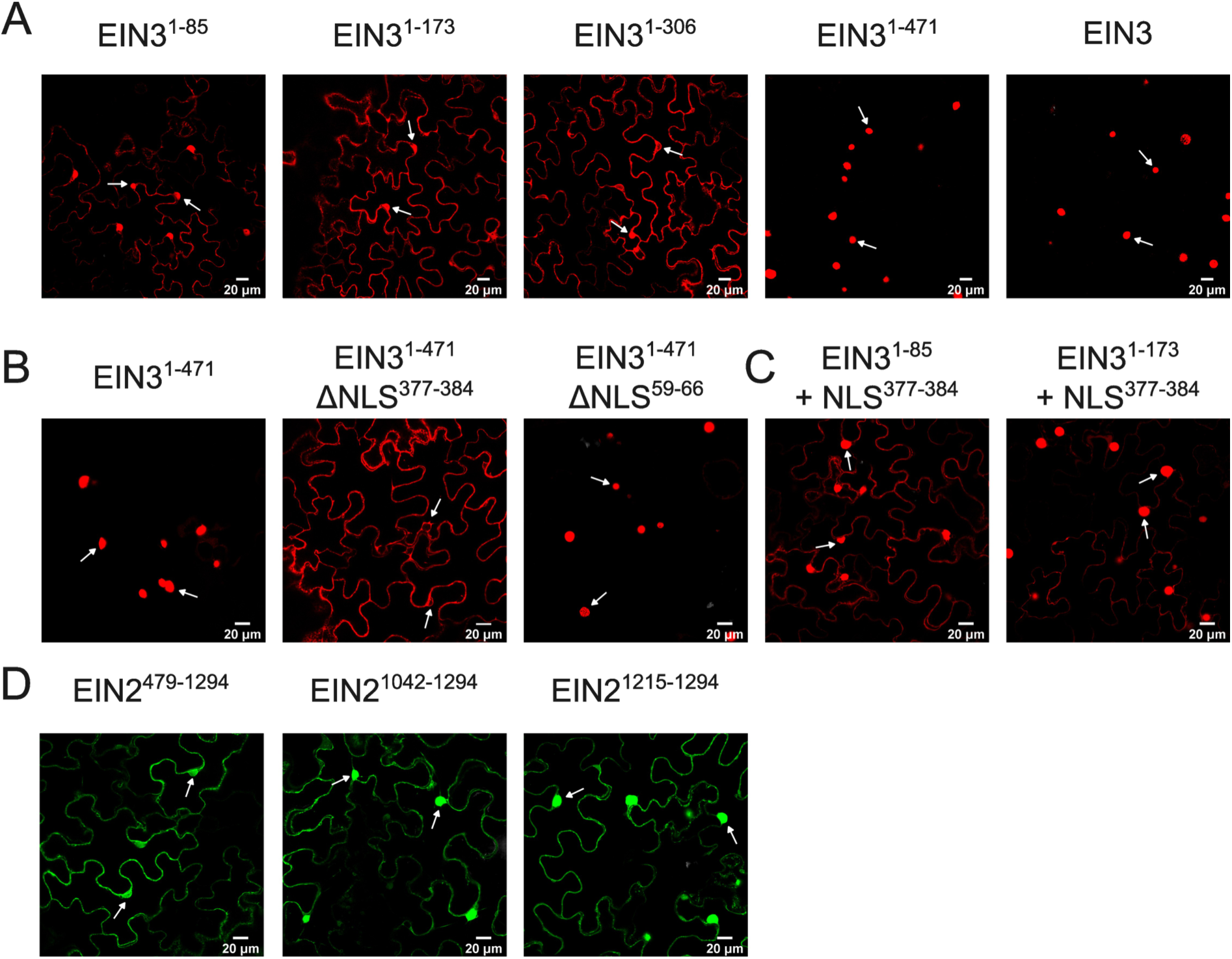
*In planta* nuclear localization of EIN2 as well as EIN3 truncations. **A)** Confocal fluorescence microscopy of EIN3 truncations fused to mCherry in transiently transformed *N. benthamiana* epidermal cells. **B)** Localization of EIN3^1–471^ and two NLS-deletion variants fused to mCherry **C)** EIN3^1–85^ and EIN3^1–173^ fused to the C-terminal NLS (residues 377-384) and mCherry **D)** Confocal fluorescence microscopy of EIN2 truncations fused to GFP. In all panels, white arrows indicate nuclei and scale bars represent 20 µm.

**Figure 7:**
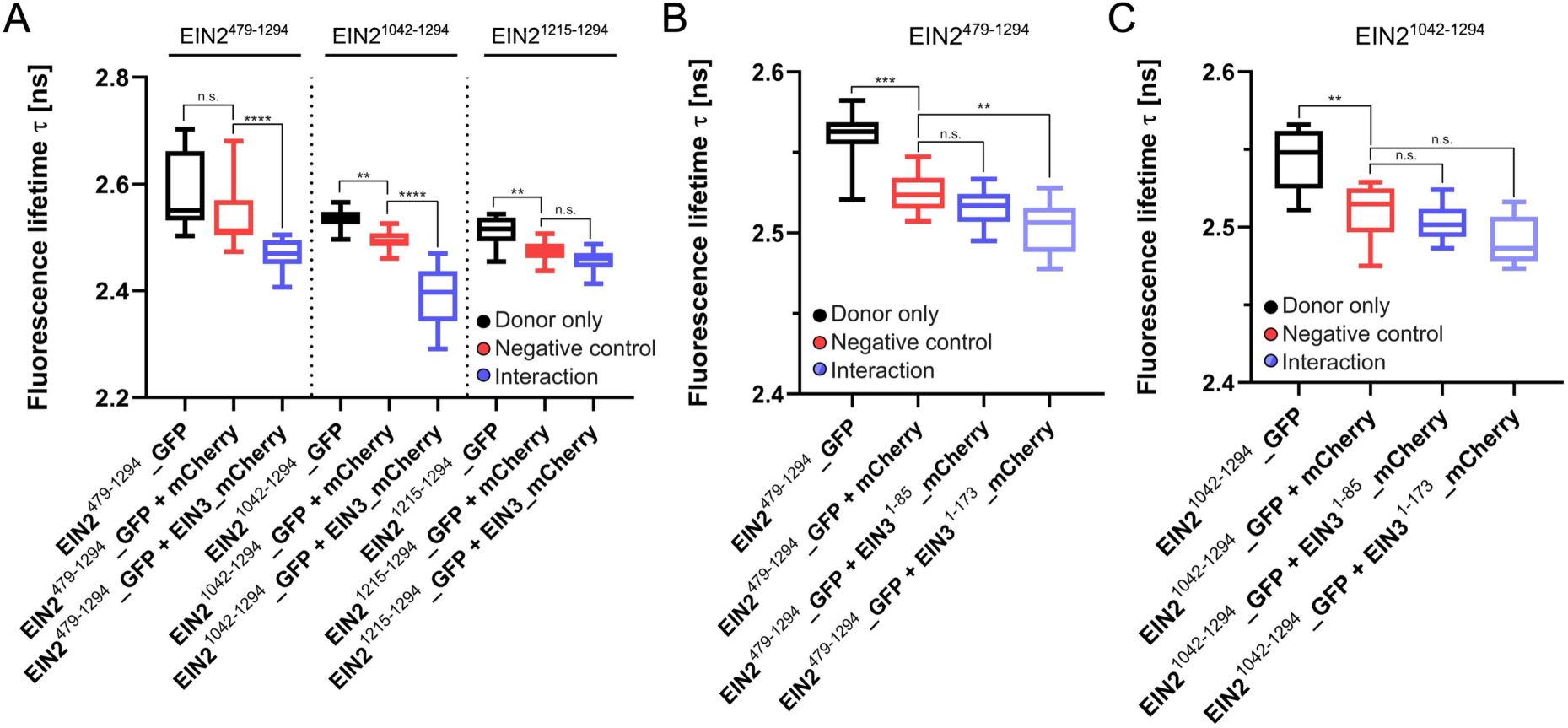
*In planta* FLIM-FRET analysis of EIN2-EIN3 interactions in *N. benthamiana*. **A)** FLIM-FRET measurements using EIN2^479–1294^, EIN2^1042–1294^, and EIN2^1215–1294^ fused to GFP as donors and full-length EIN3 fused to mCherry as acceptor. **B)** FLIM-FRET experiments using EIN2^479–1294–GFP^ as donor and EIN3^1–85^-mCherry and EIN3^1–173^–mCherry as acceptors. **C)** FLIM-FRET experiments using EIN2^1042–1294^ fused to GFP and EIN3^1–85^-mCherry or EIN3^1–173^–mCherry as acceptors. In all experiments, EIN2 truncations fused to GFP served as donors and EIN3 constructs fused to mCherry as acceptors. Donor-only measurements (black) were included for lifetime normalization, and mCherry fused to an NLS (mCherry-NLS) served as a negative control (red) to account for bystander FRET. The tested EIN2-EIN3 FRET pairs are depicted in blue. Due to weak nuclear localization, EIN3^1–85^ and EIN3^1–173^ were fused to the internal EIN3-NLS identified in Figure 6. Boxplots show the median (line), 25-75th percentile (box) and 10-90th percentile (whiskers) from pooled datasets comprising at least 16 measurements per condition. Values outside the 10-90th percentile are omitted for clarity. Statistical analysis was performed using Kruskal-Wallis and Dunn’s multiple comparisons test. *P-value <0.05; **P-value <0.01; ***P-value <0.001; ****P-value <0.0001 (donor only vs. interaction // donor only vs. negative control // negative control vs. interaction).

Because EIN3 is a transcription factor, BD-EIN3 auto-activation was also examined (Figure 5C). Full-length EIN3 showed strong reporter activation that was not suppressed by 3-AT, whereas BD-EIN3^1–306^ lacked auto-activation, consistent with previous reports^26^. Extension to BD-EIN3^1–471^ restored moderate reporter activity, which was fully suppressed by 3-AT, indicating that residues 307-471 contribute to transcriptional activation, but additional C-terminal regions are required for full activity. To refine the EIN3 dimerization domain, BD-EIN3^1–306^ was tested against a panel of AD-EIN3 truncations (Figure 5D). Interaction was detected with AD-EIN3^1–306^ and AD-EIN3^1–173^, but not with AD-EIN3^1–85^. These results narrow the EIN3 dimerization domain to residues 113-173, consistent with earlier studies but providing a more precise boundary.

### *In planta* FLIM-FRET analysis reveals the EIN2-EIN3 interaction and defines the nuclear localization requirements of both proteins

To assess the EIN2-EIN3 interaction in a physiological context, we turned to *in planta* FLIM-FRET measurements following transient expression of fluorescently tagged protein truncations in *N. benthamiana*. Because EIN2-EIN3 binding must occur in the nucleus, we first examined the subcellular localization of all constructs. EIN2 truncations fused to GFP localized to both cytoplasm and nucleus consistent with previous reports for EIN2^479–1294^ ^7,9,21^. Shorter EIN2 variants, EIN2^1042–1294^ and EIN2^1215–1294^, showed progressively stronger nuclear accumulation, reflecting the position of the EIN2 nuclear localization signal (NLS) at residues 1262-1269. EIN2^1215–1294^ also showed pronounced nucleolar enrichment, likely due to size-dependent trafficking rather than physiological relevance. In contrast, full-length EIN3 and EIN3^1–471^ localized exclusively in the nucleus, whereas shorter EIN3 truncations (EIN3^1–306^, EIN3^1–173^ and EIN3^1–85^) showed markedly reduced nuclear accumulation. Sequence analysis and PredictNLS ^28^ identified two putative NLS motifs at residues 59-66 (N-terminal NLS) and 377-384 (C-terminal NLS). Deletion analysis demonstrated that the C-terminal NLS (residues 377-384) is essential for the nuclear import, while the N-terminal NLS contributes little or not at all.

To enable FLIM-FRET measurements, we restored nuclear localization of EIN3 truncations by fusing the identified C-terminal NLS (MRKRKPNR) to their C-termini, which substantially increased nuclear-localized fluorescence signal and confirmed the functional relevance of this motif. FLIM-FRET measurements were performed using EIN2 truncations fused to GFP (donor) and EIN3 constructs fused to mCherry (acceptor). Donor only controls were included for lifetime normalization and mCherry fused to an NLS (mCherry-NLS) served as a negative control for bystander FRET. All EIN2-EIN3 combinations showed reduced donor lifetimes relative to donor-only controls. However, only EIN2^479–1294^–GFP and EIN2^1042–1294^–GFP exhibited significantly shorter lifetimes than their respective negative controls. EIN2^1215–1294^–GFP did not differ from the negative control, indicating absence of specific interaction. Negative controls displayed fluorescence lifetimes ranging from 2.47 ± 0.02 ns for EIN2^1215–1294^–GFP to 2.49 and 2.50 ± 0.02 ns for the other two EIN2-GFP truncations. EIN2^479–1294^–GFP and EIN2^1042–1294^–GFP co-expressed with EIN3-mCherry showed lifetimes of 2.46 ± 0.04 ns and 2.38 ± 0.07 ns, respectively, whereas EIN2^1215–1294^–GFP exhibited a fluorescence lifetime of 2.46 ± 0.03 ns. These data demonstrate that EIN2 residues 1042-1214 are sufficient to mediate EIN3 binding *in planta*, while residues beyond 1214, including the EIN2 NLS, are not required. To further define the EIN3 interaction surface, we tested EIN3^1–85^-mCherry and EIN3^1–173^–mCherry as acceptors. Co-expression of EIN2^479–1294^–GFP with EIN3^1–173^–mCherry significantly reduced donor lifetimes relative to the negative control, whereas EIN3^1–85^-mCherry did not. Fluorescence lifetimes of EIN2^479–1294^–GFP were 2.52 ± 0.01 ns with the negative control, 2.52 ± 0.01 ns with EIN3^1–85^-mCherry and 2.50 ± 0.02 ns when co-expressed with EIN3^1–173^–mCherry. When EIN2^1042–1294^–GFP served as the donor, neither EIN3 truncation produced significant lifetime reductions, indicating that the distal EIN2 region interacts with the C-terminal portion of EIN3 beyond residue 173. These results confirm that EIN3 residues 86-173 constitute the core EIN2 binding domain and that the distal EIN2 region engages a separate C-terminal surface of EIN3.

Together, the FLIM-FRET results corroborate the *in vitro* MST and Y2H data and define the minimal EIN2 region capable of engaging EIN3 in a physiological environment.

### Structural modeling, docking and sequence conservation analysis identify a putative EIN3 and EIN2 binding interface

To further refine the molecular interface underlying the EIN2-EIN3 interaction, we generated homology models of *Arabidopsis thaliana* EIN2 (residues 479–995) and EIN3 (full length) and performed protein-protein docking using HADDOCK^29^ (Figure 8A-D). Docking results were ranked according to the HADDOCK score (Cluster size, Van der Waals forces, electrostatic and desolvation energy for each docking cluster are shown in supplementary tables S1-S3), where positive values indicate unfavorable binding and negative values reflect favorable interactions. The EIN2 model was systematically docked against the experimentally defined EIN3 interaction region (residues 86–173). Among all tested regions, EIN2^646–674^ produced the most favorable HADDOCK score (−40), indicating a preferred interaction surface (Figure 8A). To validate this result, we performed reciprocal docking using EIN2^646–674^ against sequential EIN3 regions. A stepwise evaluation from the N- to the C-terminus revealed that the N-terminal portion of EIN3 (up to residue 200), contributes most strongly to the binding. The most favorable docking scores were obtained for EIN3^75–100^ with (−90) and EIN3^100–150^ (−75) (Figure 8B). Fine-mapping, using overlapping EIN3 segments spanning residues 86 to 215 narrowed the predicted interaction interface to EIN3 residues 86 to 100 (Figure 8C), consistent with the MST, FLIM-FRET and Y2H data identifying residues 86-173 as essential for binding. This putative EIN2 binding surface is in close proximity to the DNA binding domain (DBD) (residues 174-306) of EIN3 (Figure 8D-E).

**Figure 8:**
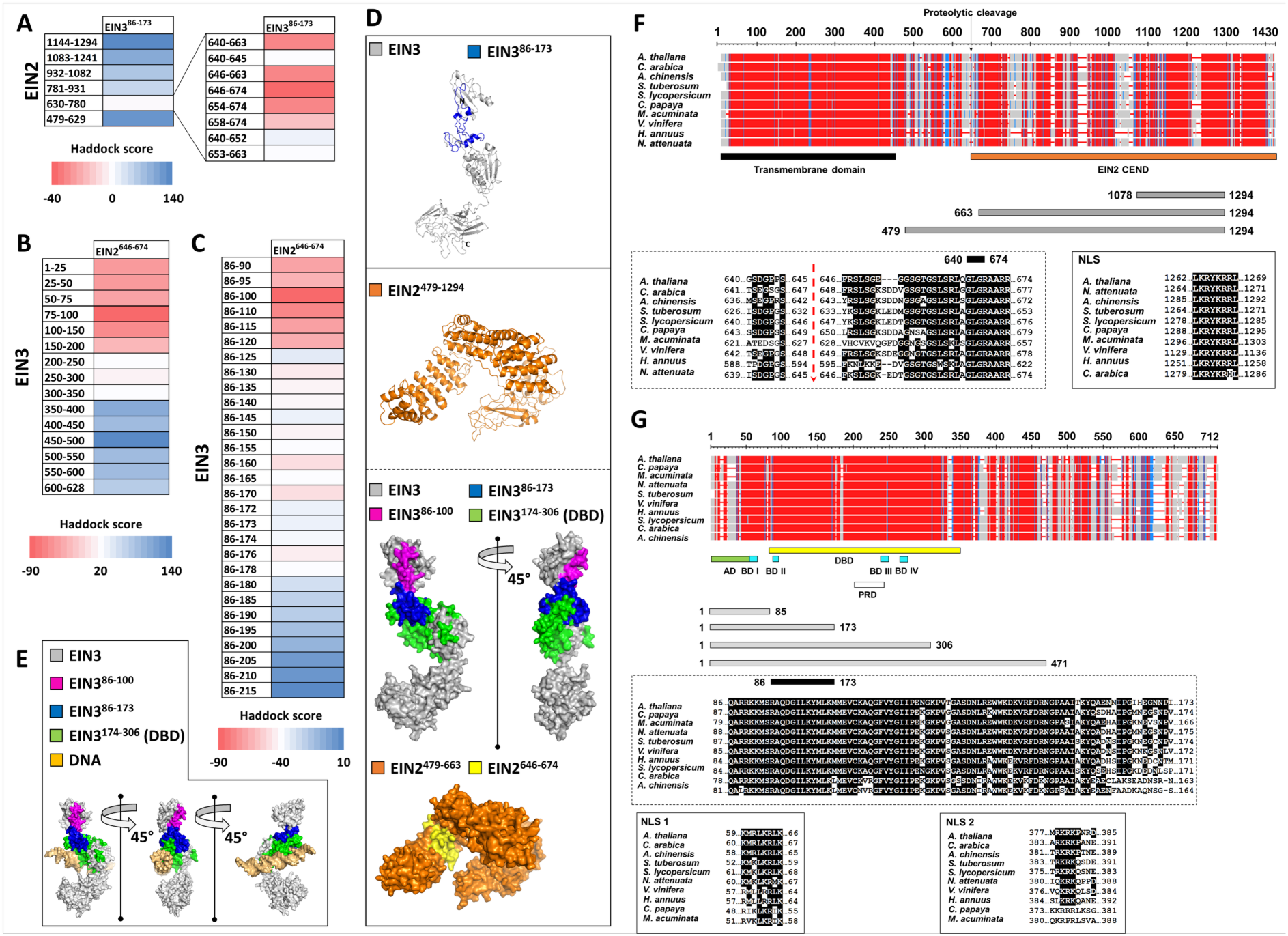
Structural modeling, docking, and sequence conservation of EIN2 and EIN3. **A)** Docking analysis of predicted interaction interfaces between EIN3^86–173^ and EIN2 truncations. Structural models were generated using I-TASSER ^37^ and are shown in ribbon and surface representation. EIN3^86–173^ is highlighted in blue, whereas EIN2^479–663^ and its truncated variants are shown in orange. Docking results were ranked by HADDOCK score, with more negative values indicating more favorable interactions. **B)** Reciprocal docking of EIN2^464–674^ against sequential EIN3 segments. **C)** Fine-mapping of the interaction surface using overlapping EIN3 fragments (residues 86-215) docked against EIN2^464–674^. **D)** Representative structural models of *Arabidopsis thaliana* EIN3 and *At*EIN2^479–1294^. The EIN3 core binding regions (residues 86-173 and 86-100) are highlighted in blue and pink, respectively, while the DNA binding domain (residues 174-206) is marked in green. EIN2 models are depicted in orange with the identified core binding region (residues 646-674) highlighted in yellow. **E)** Predicted EIN3-DNA complex (Haddock score of −37). DNA is shown in yellow, whereas EIN3 fragments follow the color scheme in panel D). **F) and G)** Multiple sequence alignment of EIN2 (B) and EIN3 (C) orthologs from ten angiosperm species (*A. thaliana*, *C. arabica*, *A. chinensis*, *S. tuberosum*, *S. lycopersicum*, *C. papaya*, *M. acuminata*, *V. vinifera*, *H. annuus* and *N. attenuate)*. Alignments include conservation plots, annotated functional domains, predicted binding regions, protein constructs used in this study, and nuclear localization signals (NLSs). Conserved regions corresponding to the predicted EIN2-EIN3 interaction interface are highlighted.

To assess the evolutionary conservation of the predicted interface, we performed multiple sequence alignments of EIN2 and EIN3 orthologs from *A. thaliana*, *C. arabica*, *A. chinensis*, *S. tuberosum*, *S. lycopersicum*, *C. papaya*, *M. acuminata*, *V. vinifera*, *H. annuus* and *N. attenuata* using T-Coffee^30^ (see Figure 8F-G). The selected species represent a wide evolutionary spectrum of angiosperms. This diversity enables robust assessment of evolutionary conservation and divergence of EIN2 and EIN3 across distinct plant lineages. EIN2 displayed strong conservation across species, with the N-terminal transmembrane region being most conserved (Figure 8F). The EIN2 constructs used in this study derived from the soluble C-terminal domain (residues 479–1294). Notably, the docking-identified region EIN2^646–674^ corresponding to the N-terminus of the EIN2-C fragment after cleavage at Ser645/Phe646 ^8^ is highly conserved across species, with the exception of *Musa acuminata* (Figure 8F). The C-terminal NLS (1262-LKRYKRRL-1269) is also strongly conserved.

EIN3 exhibited high conservation in its N-terminal region, which contains several functional elements, including the acidic domain (AD), proline-rich domain (PRD), four basic domains (BD I–IV), and the DNA-binding domain (DBD). In contrast, the C-terminal region showed substantial divergence (Figure 8G). The EIN3 constructs analyzed here predominantly encompass the conserved N-terminal portion. Within the experimentally identified interaction region, residues 86–153 were strongly conserved, whereas residues 154–173 showed greater variability (Figure 8G). Sequence analysis also revealed that the N-terminal NLS (residues 59-66, KMRLKRLK) is highly conserved, while the C-terminal NLS (residues 377-384, MRKRKPNR) is conserved mainly in its core RKRK motif (Figure 8G). Given that only the C-terminal NLS is required for nuclear localization, the strong conservation of the N-terminal NLS suggests a potential auxiliary or backup role.

Together, structural modeling, docking and evolutionary analysis converge on a conserved EIN2-EIN3 interaction interface involving EIN2 residues 646-674 and EIN3 residues 86-120, providing mechanistic support for the experimentally defined binding regions.

## DISCUSSION

Despite substantial progress in elucidating ethylene signaling, the molecular events linking EIN2 nuclear translocation to transcriptional activation remain incompletely defined. By integrating quantitative biophysics, yeast genetics, *in planta* imaging, and structural modeling, we demonstrate that EIN2 directly engages EIN3 and delineate the minimal regions required for this interaction. Together, these complementary approaches reveal a modular interaction landscape in which distinct EIN2 surfaces contribute differentially to EIN3 binding depending on affinity and cellular context. Our MST and docking analyses converge on EIN2 residues 479-662 as the primary high-affinity interaction hotspot. This region is evolutionarily conserved, yields the most favorable docking scores, and dominates binding in controlled biophysical assays. Correspondingly, EIN3 constructs containing residues 86-173 display the highest affinity for EIN2, and further C-terminal extension does not enhance binding, identifying EIN3^86–173^ as the core interaction region. MST experiments also revealed pronounced changes in initial fluorescence (IF) when EIN2 truncations served as fluorophores. Control experiments exclude dye artifacts (Figure S1), suggesting that EIN3 binding induces conformational rearrangements within the intrinsically disordered EIN2 C-terminus. ENAP1, previously proposed to stabilize an EIN2-EIN3 complex, bound both proteins with appreciable affinity (Figure 3B), consistent with its proposed chromatin-associated role^15^, yet MST-based competition assays (Figure 3B) show that ENAP1 and EIN3 compete for the same EIN2 region at elevated ENAP1 concentrations. Thus, ENAP1 does not universally stabilize an EIN2-EIN3 complex but instead acts in a dynamic, concentration-dependent manner that can either facilitate or antagonize EIN3 recruitment. The fact that ENAP1 and EIN3^1–173^ engage the same EIN2 segment further supports this interpretation and argues against a constitutive ternary complex.

Yeast two-hybrid (Y2H) assays provided additional support for a direct EIN2-EIN3 interaction, and aligned well with the intrinsic affinity landscape defined by MST despite overall weaker reporter activation in yeast due to crowding and folding constraints of the system^31^. Co-expression of EIN3 constructs consistently enhanced reporter activity above the auto-activation background of BD-EIN2^479–1294^, and the observation that EIN3^1–173^ but not EIN3^1–85^ supported interaction reinforces the MST-derived conclusion that residues 86-173 constitute the core EIN2-interaction region. Y2H analysis also clarified the functional organization of EIN3, showing that full-length EIN3 displayed strong auto-activation consistent with earlier studies^26^. EIN3^1–471^ showed intermediate activity, and only constructs containing the C-terminal region beyond residue 471 retained full transcriptional activation capacity, indicating that activation, DNA binding, and dimerization reside in distinct domains. The failure of BD-EIN2^663–1294^ to interact with EIN3 in yeast, despite comparable affinities *in vitro*, likely reflects differences in folding, stability or nuclear accessibility rather than absence of binding.

To assess interaction under more physiological conditions, we performed *in planta* FLIM-FRET measurements, ensuring nuclear accumulation of EIN3 by fusing its essential C-terminal NLS (residues 377-384) to truncations lacking it. EIN2 fragments showed progressively stronger nuclear localization with decreasing size, consistent with passive diffusion of proteins <60 kDa through the nuclear pore complex ^32^, and the pronounced nucleolar accumulation of EIN2^1215–1294^ likely reflects this size-dependent trafficking rather than physiological relevance, as this fragment is not known to occur *in vivo*. FLIM-FRET analyses revealed robust lifetime reductions for EIN2^479–1294^ and EIN2^1042–1294^ when co-expressed with EIN3, whereas EIN2^1215–1294^ showed no interaction, demonstrating that residues 1042-1214 are sufficient to mediate EIN3 binding *in planta*. These findings align with MST data showing that EIN2^1078–1294^ retains measurable, albeit lower affinity for EIN3 (low µM affinity instead of nM affinity), and explain why Y2H assays fail to detect interaction with shorter EIN2 fragments whose binding strength likely falls below the detection threshold of the yeast system.

Complementary FLIM-FRET experiments with EIN2^1042–1294^ and EIN3 truncations ruled out involvement of the EIN3 core binding domain (amino acids 86-173) in this distal interaction. Importantly, the ability of EIN2^1042–1294^ to interact with EIN3 *in planta* reflects the transient overexpression context of agroinfiltration, where elevated nuclear protein concentrations, and molecular crowding stabilize weak or transient contacts. Under such non-physiological conditions, the contribution of the high-affinity region is masked by mass-action effects, and detectable interaction does not necessarily indicate sufficiency under endogenous expression levels. This distinction is underscored by the classical rescue experiments in *ein2* mutants. Only EIN2 fragments that include the high-affinity region such as 454/459/479-1294 or 645-1294 fully restore ethylene responses^5,8,9,33^. These fragments retain either the complete or the C-terminal portion of the primary hotspot together with the distal interaction region, ensuring sufficient binding capacity at endogenous nuclear concentrations. In contrast, the distal region alone is unlikely to support robust EIN3 engagement under physiological expression levels.

*In silico* docking identified a highly conserved interaction hotspot within *A. thaliana* EIN2 residues 646-674 and placed the corresponding EIN3 contact surface within residues 86-100, providing a structural rationale for the high-affinity interaction detected by MST and the Y2H-defined requirement for EIN3^1–173^. This docking-derived interface aligns with the intrinsic affinity landscape revealed by MST, which consistently identified EIN2^479–662^ as the dominant high-affinity region, and it complements Y2H results showing that only EIN3 constructs containing residues 86–173 support interaction. At the same time, FLIM-FRET demonstrated that a more distal EIN2^1042–1214^ segment is sufficient to mediate detectable interaction under transient overexpression, indicating that EIN2 harbors multiple interaction-competent surfaces whose contribution depends on affinity and cellular context.

Together, these biochemical, cellular, and structural analyses support a two-tiered model in which a conserved, specificity-defining high-affinity interface (EIN2^479–662^ / EIN3^86–173^) underpins physiological signaling, while a secondary, lower-affinity surface downstream of residue ∼1042 can contribute under conditions of elevated local concentration or in the presence of stabilizing cofactors. This framework reconciles the intrinsic affinity landscape with the context-dependent behavior observed *in planta* and provides a mechanistic basis how EIN2 engages EIN3 to initiate ethylene transcriptional responses.

## MATERIAL AND METHODS

### Prediction of EIN2 and EIN3 protein-protein interactions (PPI) and functional networks in *A. thaliana*

Protein–protein and gene interaction analyses for EIN2 and EIN3 were performed using the STRING database, which integrates experimentally validated interactions, computational prediction, and text mined associations^34^. A PPI network was constructed to visualize their roles within the ethylene response pathway. A high-confidence threshold (interaction score ≥ 0.7) was applied and analysis was restricted to *Arabidopsis thaliana*, to ensure species-specific relevance. Gene co-expression information available within STRING was also incorporated. Complementary network analyses were conducted using FunCoup 6, which integrates additional evidence types such as mRNA co-expression and phylogenetic profile similarity, where conservation across species indicates functional association^24^. Default parameters were used unless otherwise stated.

### Age dependent expression patterns of EIN2 and EIN3 and related proteins in *A. thaliana*

The expression dynamics of ETR1, CTR1, EIN2, EBF1, EBF2, EIN5, UPF1, EIN3, EIL1, ENAP1, and ENAP2 were examined across a 16-day developmental time course in *Arabidopsis thaliana*. Publicly available transcriptomic datasets from the Expression Atlas repository were used for this analysis^23^. Transcript abundancies reported as transcripts per million (TPM), a normalization metric that reflects the relative abundance of transcripts for each gene or isoform, enabling quantitative comparison across samples.

### Multiple interspecies sequence alignment for EIN2 and EIN3

EIN2 and EIN3 protein sequences were retrieved from the Uniprot database (see Table S4), and multiple sequence alignments were performed using T-COFFEE^30,35^ and COBALT (Constraint-based Multiple Alignment Tool)^36^. For EIN2, the alignment focused on the major interaction region identified in this study (residues 479 to 663). For EIN3, both the full-length proteins and the multi-functional interaction region comprising residues 86 to 173 were aligned to assess conservation across species.

### Homology modelling of EIN2 and EIN3 core regions and docking experiments

Three-dimensional homology models of EIN2^479–995^ and EIN3 full length were generated using the I-TASSER protein structure prediction server^37^. Both proteins contain extensive intrinsically disordered and flexible regions, which limited the reliability of structure prediction using AlphaFold and Chai. Therefore, I-TASSER was selected as an alternative approach capable of producing homology models suitable for downstream structural and functional investigations. Model construction was based on the 10 closest homologs identified by TM-alignment. For each comparison, I-TASSER reports the optimal structural alignment between two proteins and the corresponding TM-score, where values >0.5 indicate that two proteins share the same overall fold. Model quality was further evaluated using the C-score, a confidence metric that estimates the reliability of predicted protein structures and typically ranges from −5 to 2. Higher C-scores correspond to increased confidence and improved model quality, while values greater than −1.5 are generally indicative of correct global topology. Model quality evaluation revealed high confidence, with TM-scores above 0.6, consistent with correct fold prediction (TM-score > 0.5). The C-scores were −1.31 for EIN2^479–995^ and −1.03 for EIN3, falling within the typical range of −5 to 2, indicating reliable model quality. Model construction was performed using multiple template structures. The EIN2^479–995^ model was generated based on PDB templates 5A1U, 7BII, 4A0C, 7JQ9, 2QNA, 6YEJ, 6EZ8, 6N1Z, 3EA5, and 6AHO. In parallel, the EIN3 model was built using the following PDB entries: 3CHN, 7CUN, 5U1S, 6Z2W, 7ABI, 6UM1, 2NYY, 2UVC, 6EGO, and 3CMX. Protein-protein docking simulations between EIN3-EIN2 were carried out using HADDOCK^29^. Docking poses were ranked according to the HADDOCK scoring function (corresponding binding energies and cluster size are shown in supplementary tables S1-S3), where positive scores denote unfavorable interactions and negative scores indicate energetically favorable binding. The full-length EIN3 model was used and residues in different regions were activated in the docking approach to be involved in the interaction with EIN2 (e.g., EIN3^86–173^, EIN3^75–100^, EIN3^100–150^). For EIN2, multiple truncated variants were screened including EIN2^479–995^, EIN2^479–663^, EIN2^640–663^, EIN2^640–645^, EIN2^646–663^, EIN2^646–674^, EIN2^654–674^, EIN2^658–674^, EIN2^640–652^ and EIN2^653–663^. All figures were made using the PyMOL Molecular Graphics System, Version 1.3, Schrödinger, LLC.

### Expression of EIN2 truncations

The following EIN2 constructs were successfully expressed and purified: EIN2^479–1294^, EIN2^663–1294^ and EIN2^1078–1294^. EIN2^479–1294^ was expressed from the pET15b vector containing an N-terminal deca-His-tag, whereas all other constructs were expressed from the pET21a vector which provides a C-terminal hexa-His-tag. Protein expression was carried out in the *E. coli* ArcticExpress cells grown in LB media according to the manufacturer’s instructions. Briefly, Ampicillin (100 µg/mL working concentration) and Gentamycin (20 µg/mL working concentration) were added to the pre-culture, while no antibiotic was used for the main culture. After overnight incubation of the pre-culture at 37 °C, the main culture was inoculated to an initial OD_600_ of 0.1 and grown for 3 hours at 30 °C. The temperature was then reduced to 12 °C, cultures were equilibrated for 15 minutes, and protein expression was induced with 1 mM IPTG., Cells were harvested the following day by centrifugation (7.000 x g, 10 minutes), flash-frozen in liquid nitrogen and stored at −20 °C until usage.

### Expression of EIN3 truncations

EIN3^1–85^, EIN3^1–173^, EIN3^1–306^ and EIN3^1–471^ were cloned into the pETEV16b (a modified pET16b backbone containing a TEV cleavage site between the GOI and the His-tag). EIN3^1–85^ and EIN3^1–173^ were expressed in *E. coli* BL21pRARE cells using 2YT-media, whereas EIN3^1–306^ and EIN3^1–471^ were expressed in ArcticExpress cells following the protocol described for EIN2. The pre-culture was mixed with ampicillin (100 µg/mL working concentration) and chloramphenicol (30 µg/mL working concentration) and grown overnight at 37 °C. For main cultures, 2 % ethanol and ampicillin were added prior to inoculation to an OD_600_ of 0.1. The main culture was grown at 37 °C until an OD_600_ of 0.4 was reached. Afterwards, the temperature was decreased to 16 °C and the induction with 0.5 mM IPTG was performed between an OD_600_ of 0.6 – 0.8. Cells were harvested the next day by centrifugation (7.000 x g, 10 minutes), flash-frozen in liquid nitrogen and stored at −20 °C until usage.

### Expression of ENAP1

ENAP1 was cloned into the pETEV16b vector containing an N-terminal deca-His-tag. Expression was performed in *E. coli* BL21pRARE cells following the protocol used for EIN3 truncations, except that the expression was performed at 37 °C and cells were harvested 2 hours after IPTG induction.

### Protein purification

All proteins were purified by immobilized affinity chromatography (IMAC) essentially as described in ^22^. Briefly, the stored cell pellet was resuspended in 1× PBS, 5 % (w/v) glycerol and 0.1 % (v/v) Nonidet-P40 pH 7.4 (5 mL per g cell). DNase and antifoam were added, and cells were lysed using a high-pressure cell disruptor afterwards. After cell disruption, cell debris and insoluble protein were separated by centrifugation at 100.000 x g for 1 h, and the supernatant was applied to a pre-equilibrated Ni^2+^-IMAC column. For proteins expressed in BL21pRARE, the column was washed with 500 mM NaCl and 50 mM sodium phosphate (pH 7.4) until baseline stabilization. Non-specific proteins were removed using stepwise washes containing 50 mM and 100 mM imidazole. Target proteins were eluted with 500 mM imidazole in 500 mM NaCl, 50 mM sodium phosphate buffer (pH 7.4). For proteins expressed in ArcticExpress cells, an additional wash with 20 mM TRIS and 2 M urea (pH 6.8) was included prior to imidazole elution. Purified proteins were concentrated using ultrafiltration devices, flash-frozen in liquid nitrogen, and stored at −80 °C.

### Labeling of *At*EIN2 truncations with Alexa488-NHS ester

Purified and concentrated EIN2 proteins were first rebuffered into labeling buffer (300 mM NaCl and 50 mM NaP_i_) using a PD-10 desalting column. Alexa488-NHS ester was added in a 2.5-fold molar excess and incubated with the protein for 45 minutes at room temperature under gentle rotation in the dark. To remove unreacted dye, the reaction mixture was rebuffered into labeling buffer supplemented with 5 % glycerol (w/v) using a PD-mini desalting column. The labeled protein was concentrated using an ultrafiltration unit and centrifuged at 229.600 x g (50.000 rpm) for 35 minutes to remove aggregates. Protein and fluorophore concentration were determined using a NanoDrop One spectrophotometer. Labeling efficiency was calculated by dividing the fluorophore concentration by the protein concentration and multiplying the result with 100 %.

### Microscale Thermophoresis (MST) measurements

MST measurements were performed using a Monolith NT.115 (NanoTemper). LED power was adjusted according to labeling efficiency, while the MST power was kept constant at 40 %. For all measurements, MST Buffer (300 mM NaCl, 50 mM HEPES and 0.05 % Tween-20, pH 7.5) and standard capillaries were used. Fo assays involving *At*EIN3^1–306^, the buffer was adjusted to 8.0 due to the pI of the protein. Unless stated otherwise, data were analyzed using the Initial Fluorescence method and fitted in Origin using a logistic model. Fluorophore-labelled proteins were used at 100 nM, and ligands were titrated up to 50 µM. To this end, a serial dilution of the ligand was prepared in PCR tubes and mixed in a 1:1 ratio with 200 nM of the respective fluorophore (resulting in 100 nM fluorophore final concentration). Samples were incubated on ice for 18 min, centrifuged at 13.400 x g for 2 min, loaded into the capillaries and measured. All MST data represent three independent replicates. Negative controls include BSA as a non-binding ligand or free dye without labelled protein.

### Yeast Two-Hybrid Assay

Yeast Two-Hybrid (Y2H) Assays were performed using *S. cerevisiae* strain AH109 with pACT2 (AD-Gal4) and pGBKT7 (BD-Gal4) vectors. Constructs were cloned into either the pACT2 (Ampicillin resistance) or pGBKT7 (Kanamycin resistance) vector, the constructs were previously cloned into pDONR207 (Gentamycin resistance) which was used as the donor vector for the subsequent Gateway Cloning method (Invitrogen). The strain was kindly provided by Prof. Dr. Wilde (University of Freiburg), and empty vectors and the positive control were kindly provided by Prof. Dr. Bauer (Heinrich Heine University Düsseldorf). Co-transformations were performed via the lithium acetate (LiAc) method according to^38^. Tansformants were selected on synthetic defined (SD) medium containing adenine and all amino acids except Leu (pACT2) and Trp (pGBKT7) and incubated for 2-3 days at 30 °C. For interaction assays, overnight liquid cultures grown in SD-LW medium (lacking Leu and Trp) were adjusted to an OD_600_ of 0.1 (10^−1^) and serial dilutions down to 10^−4^ were prepared. Ten microliters of each dilution were spotted onto SD-LW plates (transformation control) and SD-LWH plates (lacking Leu, Trp and His) to test for protein-protein interaction. To increase selectivity and ensure reliable and valid interaction data, SD-LWH plates were supplemented with 0.5 mM 3-amino-1,2,4-triazole.

### Transient transformation and expression in *N. benthamiana*

Transient transformation of *N. benthamiana* abaxial leaf epidermal cells was conducted as described previously^22^ with minor changes. OD_600_ was always set to 0.5 and since agrobacterium strain GV3101:p19 was used for all expression vectors (except mCherry-NLS), the strain did not need to be added in addition. Plants were kept under a natural day/night cycle before microscopy analysis.

### Fluorescence lifetime imaging and fitting of fluorescence decays

Fluorescence lifetime imaging was performed on a Leica Stellaris 8 FALCON microscope or a ZEISS LSM780 with LSM Upgrade Kit from Picoquant. Imaging on the LSM 780 was done as described in ^39^. The fluorescence decays from data acquired using the LSM 780 were fitted using a previously reported method with a minor modification^22^.

On the Stellaris 8 FALCON (FALCON extension), a Leica HC PL APO 40x/1,1 W motCORR CS2 objective was used. GFP(S65T) and mCherry were excited with a pulsed white-light laser (40 MHz), filtered for 488 nm and 561 nm using an acousto-optical tunable filter (AOTF). For emission detection, HyD X detectors were used, and the emitted light was filtered through an acousto-optical beam splitter (AOBS) to detect a range of 493-534 nm (GFP) and 590-636 nm (mCherry). Images were acquired at zoom 10 with a resolution of 256×256 pixels (each pixel corresponding to an area of 0.01299 µm² (113.97 × 113.97 nm)), and a pixel dwell time of 15.24 µs. The number of frames was slightly adjusted to reach comparable photon counts per pixel in each measurement. Nuclei containing high donor concentrations were avoided to prevent pile-up effects.

Fluorescence decays of selected ROIs in the FLIM images were analyzed within Leica Application Suite X (LAS X, version 4.8.2). Decays were fitted using the inbuilt FLIM fitting tool (Fitting model: n-exponential reconvolution). For donor-only samples one lifetime was fitted (model parameter n = 1). For all other samples, one lifetime was fixed to the arithmetic mean of the lifetime of the donor-only samples, and a second lifetime was fitted within the limits of 10 % and 80 % of this value (model parameter n = 2). The intensity weighted lifetime was regarded as the apparent lifetime of the sample. FRET efficiency was calculated from the intensity weighted lifetime of the FRET sample over the arithmetic mean of the intensity weighted lifetimes of the donor-only samples.

### Fluorescence imaging

Images showing protein localization in *N. benthamiana* were acquired with a ZEISS LSM 780 or a Leica Stellaris 8 microscope using the same objectives as described for FRET-FLIM experiments. GFP and mCherry were excited at 488 nm or 561 nm, respectively. Signal was acquired around the maximum emission peaks of the fluorophores. Laser power was adjusted differently based on signal intensity.

### Statistical analysis

For statistical analysis, GraphPad Prism 8 was used. For the *in planta* FLIM-FRET experiments, Kruskal–Wallis test combined with Dunn’s multiple comparisons test were performed.

## Data availability statement

All relevant data are included in this manuscript and its supporting information. Imaging datasets supporting the findings of this study are available in the BioImage Archive under accession S-BIAD3653. Further inquiries or requests for materials can be directed to.

## Acknowledgements

We would like to thank the Bauer’s Lab (Heinrich-Heine-University, Institute of Botany) for kindly supplying the positive control used in the Y2H assay as well as the Wilde’s Lab (University of Freiburg, Institute for Molecular Genetics of Prokaryotes) for providing the AH109 strain.

## Funding

This work was supported by a grant of the Deutsche Forschungsgemeinschaft (DFG, German Research Foundation) to G.G. (project ID 504365166). We would like to acknowledge the Center for Advanced Imaging (CAi) at Heinrich-Heine-University Düsseldorf for providing access to to the Zeiss LSM 780 + 4-channel FLIM exten. (Picoquant), funded by the Deutsche Forschungsgemeinschaft (DFG, German Research Foundation) – Project-ID DFG-INST 208/551-1 FUGG as well as the Leica STELLARIS 8, funded by the Deutsche Forschungsgemeinschaft (DFG, German Research Foundation) – Project-ID INST 208/941-1 FUGG. Meik Thiele funded by the Deutsche Forschungsgemeinschaft (DFG, German Research Foundation) under Germanýs Excellence Strategy – EXC-2048 – project ID 390686111 and Jan Eric Maika was supported by the Deutsche Forschungsgemeinschaft (DFG, German Research Foundation) through CSCS (FOR5235).

## Author contributions

Conceptualization, G.G and R.S; Methodology, F.W., M.H. and G.G.; Investigation, F.W., M.T. and J.E.M.; Formal analysis, F.W. and M.T.; Validation, F.W. and M.T.; Visualization, F.W. and R.J.E.; Writing – Original draft, F.W., M.T., R.J.E., R.S. and G.G.

All authors read and edited the manuscript prior to publication.

## Competing Interests statement

The authors declare that they have no known competing financial interests or personal relationships that could have appeared to influence the work reported in this paper.

## ABBREVIATIONS

AD: Activation domain
At: *Arabidopsis thaliana*
BD: Binding domain
CTR1: CONSTITUTIVE TRIPLE RESPONSE1
DBD: DNA-binding domain
EBF1/2: EIN3 BINDING F-BOX PROTEINS ½
EBS: EIN3 binding site
ER: endoplasmic reticulum
EIN2-5: ETHYLENE INSENSITIVE2-5
EIN2-C: C-terminal domain of EIN2
EIL1-5: EIN3-like 1-5
ENAP1/2: EIN2 NUCLEAR ASSOCIATED PROTEIN 1/2
ETR: ethylene receptor
FLIM-FRET: fluorescence lifetime microscopy and Förster’s resonance energy transfer
GAL4: galactose-responsive transcription factor
NLS: Nuclear localization sequence
NRMAP: natural resistance-associated macrophage protein
SD-agar: synthetic defined agar
UPFs: up-frameshift proteins or nonsense-mediated decay proteins
Y2H: Yeast Two Hybrid

## Notes

### Competing Interest Statement

The authors have declared no competing interest.

