## Supporting Information for "Structural Mapping of the EIN2–EIN3 Interaction Core and Its Integration with ENAP1 in Ethylene Signaling"

### Table of contents

**Fig. S1.** *In vitro* MST binding analysis of representative negative controls.
**Fig. S2.** Y2H assay with positive hits of the EIN2-EIN3 interaction with different 3-AT concentrations (1, 2, and 3 mM) after 10, 12 and 14 days.
**Table S1.** Binding Energies and size of the docking clusters analyzed in Figure 8A.
**Table S2.** Binding Energies and size of the docking clusters analyzed in Figure 8B.
**Table S3.** Binding Energies and size of the docking clusters analyzed in Figure 8C.
**Table S4.** Information of the EIN2 and EIN3 sequences used in the multiple sequence alignment.

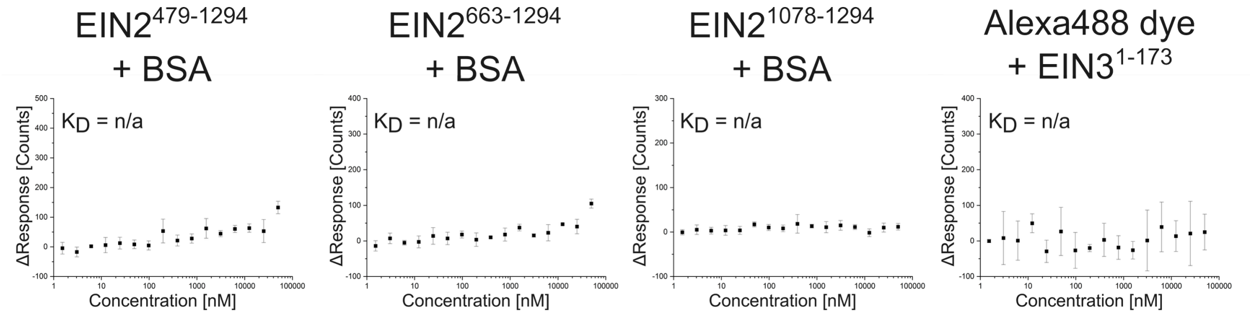

**Figure S1: *In vitro* MST binding analysis of representative negative controls.** Binding of labeled EIN2 truncations was tested against BSA to check for unspecific interaction. To eliminate unspecific dye interactions, EIN3$^{1\text{-}173}$ was used as a representative and titrated against the Alexa488 dye. MST binding data were analyzed via Initial Fluorescence analysis and curves were fitted using the Logistic Fit function in Origin. Data points represent mean values and standard deviation based on three independent measurements (n = 3).

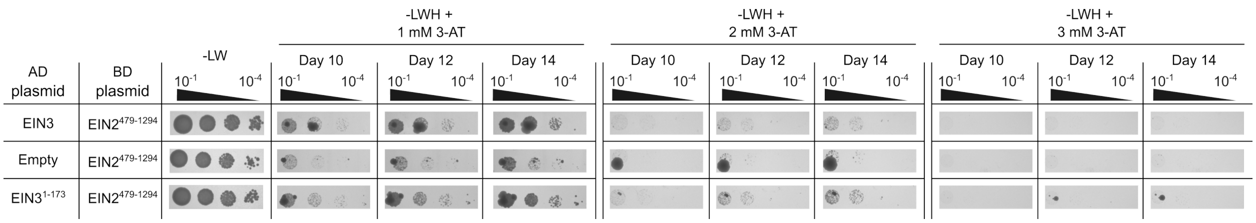

**Figure S2: Y2H assay with positive hits of the EIN2-EIN3 interaction with different 3-AT concentrations (1, 2, and 3 mM) after 10, 12 and 14 days.** EIN2$^{479\text{-}1294}$ was fused to the binding domain of GAL4 (BD plasmid), whereas EIN3 truncations were fused to the activation domain of GAL4 (AD plasmid). Yeast strain AH109 co-transformed with both plasmids was spotted in a 10-fold serial dilution ($OD_{600}$ = $10^{-1}$ – $10^{-4}$) on synthetic defined (SD) agar plates lacking leucine and tryptophane (-LW) (transformation control) as well as lacking leucine, tryptophane and histidine (-LWH) (selection for protein interaction). SD -LWH agar plates additionally contained either 1, 2 or 3 mM 3-amino-1,2,4-triazole (3-AT) for better selectivity. Images of SD -LW plates were taken after three days, while -LWH plates were imaged after 10,12, and 14 days.

**Table S1. Binding Energies and size of the docking clusters analyzed in Figure 8A.**

| | Energy | | | | |
|---|---|---|---|---|---|
| Complex | Van der walls [kcal/mol] | Electrostatic [kcal/mol] | Desolvation [kcal/mol] | Cluster size | Score |
| $EIN2^{479-629}$-$EIN3^{86-173}$ | -29.7 ± 4.2 | -373.8 ± 58.2 | 30.7 ± 7.0 | 8 | 110 |
| $EIN2^{630-780}$-$EIN3^{86-173}$ | -44.0 ± 18.0 | -123.9 ± 37.9 | -23.1 ± 1.5 | 10 | 9 |
| $EIN2^{781-931}$-$EIN3^{86-173}$ | -39.7 ± 3.9 | -166.8 ± 54.8 | -7.6 ± 3.4 | 11 | 47 |
| $EIN2^{932-1082}$-$EIN3^{86-173}$ | -32.7 ± 3.7 | -393.7 ± 112.8 | 2.8 ± 1.7 | 7 | 63 |
| $EIN2^{1083-1241}$-$EIN3^{86-173}$ | -27.7 ± 3.3 | -290.9 ± 48.9 | 17.1 ± 2.5 | 7 | 90 |
| $EIN2^{1144-1294}$-$EIN3^{86-173}$ | -21.6 ± 2.9 | -389.4 ± 73.7 | 26.9 ± 3.4 | 6 | 120 |
| $EIN2^{640-663}$-$EIN3^{86-173}$ | -29.7 ± 4.8 | -213.3 ± 28.2 | -2.4 ± 0.9 | 26 | -30 |
| $EIN2^{640-645}$-$EIN3^{86-173}$ | -20.5 ± 3.4 | -91.7 ± 25.3 | 8.0 ± 1.4 | 17 | 10 |
| $EIN2^{646-663}$-$EIN3^{86-173}$ | -31.9 ± 4.6 | -316.0 ± 48.3 | 1.7 ± 0.3 | 41 | -30 |
| $EIN2^{646-674}$-$EIN3^{86-173}$ | -22.2 ± 9.7 | -271.3 ± 21.7 | -4.4 ± 0.9 | 14 | -40 |
| $EIN2^{654-674}$-$EIN3^{86-173}$ | -28.0 ± 7.8 | -142.3 ± 16.9 | -3.3 ± 0.5 | 14 | -30 |
| $EIN2^{658-674}$-$EIN3^{86-173}$ | -19.6 ± 7.1 | -196.3 ± 37.4 | -2.9 ± 0.9 | 25 | -10 |
| $EIN2^{640-652}$-$EIN3^{86-173}$ | -13.1 ± 6.8 | -132.7 ± 49.1 | -2.6 ± 0.6 | 16 | 20 |
| $EIN2^{653-663}$-$EIN3^{86-173}$ | -18.5 ± 6.7 | -108.7 ± 13.8 | 7.1 ± 1.2 | 19 | 10 |

**Table S2. Binding Energies and size of the docking clusters analyzed in Figure 8B.**

| | Energy | | | | |
|---|---|---|---|---|---|
| Complex | Van der Walls [kcal/mol] | Electrostatic [kcal/mol] | Desolvation [kcal/mol] | Cluster size | Score |
| $EIN3^{1-25}$-$EIN2^{646-674}$ | -29.4 ± 3.5 | -188.5 ± 9.5 | -4.8 ± 1.0 | 15 | -40 |
| $EIN3^{25-50}$-$EIN2^{646-674}$ | -25.9 ± 1.6 | -230.4 ± 9.5 | -5.5 ± 1.3 | 17 | -40 |
| $EIN3^{50-75}$-$EIN2^{646-674}$ | -27.2 ± 2.7 | -215.4 ± 20.4 | -6.1 ± 2.9 | 11 | -30 |
| $EIN3^{75-100}$-$EIN2^{646-674}$ | -38.1 ± 4.7 | -239.0 ± 20.9 | -9.2 ± 1.2 | 56 | -90 |
| $EIN3^{100-150}$-$EIN2^{646-674}$ | -31.7 ± 3.0 | -240.0 ± 15.7 | -11.5 ± 2.1 | 12 | -55 |
| $EIN3^{150-200}$-$EIN2^{646-674}$ | -37.3 ± 5.3 | -187.1 ± 15.0 | -6.2 ± 1.1 | 29 | -10 |
| $EIN3^{200-250}$-$EIN2^{646-674}$ | -15.6 ± 3.4 | -328.4 ± 44.5 | 6.9 ± 1.5 | 16 | 60 |
| $EIN3^{250-300}$-$EIN2^{646-674}$ | -19.2 ± 8.1 | -81.9 ± 39.4 | 5.6 ± 1.7 | 22 | 50 |
| $EIN3^{300-350}$-$EIN2^{646-674}$ | -17.1 ± 4.6 | -113.2 ± 28.6 | 8.4 ± 2.2 | 14 | 60 |
| $EIN3^{350-400}$-$EIN2^{646-674}$ | -27.3 ± 4.8 | -117.5 ± 41.9 | 14.9 ± 1.8 | 11 | 110 |
| $EIN3^{400-450}$-$EIN2^{646-674}$ | -33.2 ± 2.6 | -61.4 ± 14.7 | 16.6 ± 2.1 | 25 | 100 |
| $EIN3^{450-500}$-$EIN2^{646-674}$ | -24.3 ± 3.2 | -119.5 ± 24.1 | 21.6 ± 2.5 | 15 | 130 |
| $EIN3^{500-550}$-$EIN2^{646-674}$ | -26.6 ± 2.4 | -177.6 ± 31.8 | 15.3 ± 1.8 | 19 | 100 |
| $EIN3^{550-600}$-$EIN2^{646-674}$ | -22.6 ± 3.2 | -129.8 ± 34.5 | 18.3 ± 1.1 | 15 | 100 |
| $EIN3^{600-628}$-$EIN2^{646-674}$ | -25.4 ± 4.4 | -103.1 ± 33.1 | 17.5 ± 2.9 | 25 | 90 |

**Table S3. Binding Energies and size of the docking clusters analyzed in Figure 8C.**

| | Energy | | | | |
|---|---|---|---|---|---|
| Complex | Van der Walls [kcal/mol] | Electrostatic [kcal/mol] | Desolvation [kcal/mol] | Cluster size | Score |
| $EIN3^{86\text{-}90}$-$EIN2^{646\text{-}674}$ | -28.4 ± 3.8 | -246.4 ± 16.4 | -12.6 ± 2.0 | 17 | -70 |
| $EIN3^{86\text{-}95}$-$EIN2^{646\text{-}674}$ | -26.8 ± 5.0 | -202.6 ± 36.7 | -15.8 ± 1.2 | 13 | -65 |
| $EIN3^{86\text{-}100}$-$EIN2^{646\text{-}674}$ | -31.2 ± 1.5 | -255.9 ± 11.9 | -16.2 ± 2.1 | 28 | -90 |
| $EIN3^{86\text{-}110}$-$EIN2^{646\text{-}674}$ | -29.9 ± 6.5 | -249.1 ± 15.0 | -16.6 ± 1.3 | 22 | -80 |
| $EIN3^{86\text{-}115}$-$EIN2^{646\text{-}674}$ | -35.2 ± 5.1 | -124.4 ± 38.6 | -14.2 ± 3.1 | 17 | -71 |
| $EIN3^{86\text{-}120}$-$EIN2^{646\text{-}674}$ | -40.9 ± 6.6 | -80.7 ± 16.9 | -18.2 ± 0.7 | 19 | -67 |
| $EIN3^{86\text{-}125}$-$EIN2^{646\text{-}674}$ | -29.5 ± 3.3 | -204.2 ± 17.8 | -6.0 ± 2.0 | 17 | -32 |
| $EIN3^{86\text{-}130}$-$EIN2^{646\text{-}674}$ | -25.8 ± 1.7 | -179.9 ± 44.5 | -6.1 ± 1.1 | 16 | -46 |
| $EIN3^{86\text{-}135}$-$EIN2^{646\text{-}674}$ | -30.3 ± 3.3 | -276.6 ± 51.8 | -3.2 ± 1.9 | 15 | -43 |
| $EIN3^{86\text{-}140}$-$EIN2^{646\text{-}674}$ | -23.5 ± 6.6 | -200.1 ± 45.7 | -6.9 ± 1.4 | 12 | -42 |
| $EIN3^{86\text{-}145}$-$EIN2^{646\text{-}674}$ | -25.4 ± 4.9 | -195.7 ± 32.8 | -7.6 ± 2.1 | 17 | -35 |
| $EIN3^{86\text{-}150}$-$EIN2^{646\text{-}674}$ | -26.5 ± 5.5 | -220.9 ± 41.3 | -6.5 ± 2.0 | 12 | -43 |
| $EIN3^{86\text{-}155}$-$EIN2^{646\text{-}674}$ | -15.7 ± 5.3 | -240.1 ± 44.1 | -8.3 ± 3.2 | 11 | -40 |
| $EIN3^{86\text{-}160}$-$EIN2^{646\text{-}674}$ | -24.6 ± 4.5 | 194.6 ± 23.1 | -10.3 ± 3.9 | 13 | -50 |
| $EIN3^{86\text{-}165}$-$EIN2^{646\text{-}674}$ | -21.7 ± 3.4 | -216.9 ± 17.9 | -5.5 ± 2.2 | 12 | -40 |
| $EIN3^{86\text{-}170}$-$EIN2^{646\text{-}674}$ | -25.8 ± 5;2 | -190.8 ± 12.7 | -10.3 ± 2.2 | 13 | -50 |
| $EIN3^{86\text{-}172}$-$EIN2^{646\text{-}674}$ | -14.0 ± 4.2 | -227.2 ± 27.3 | -3.9 ± 1.5 | 10 | -35 |
| $EIN3^{86\text{-}173}$-$EIN2^{646\text{-}674}$ | -14.1 ± 3.9 | -226.2 ± 29.9 | -4.0 ± 1.7 | 14 | -35 |
| $EIN3^{86\text{-}174}$-$EIN2^{646\text{-}674}$ | -29.2 ± 7.1 | -201.3 ± 34.7 | -2.5 ± 0.8 | 12 | -38 |
| $EIN3^{86\text{-}176}$-$EIN2^{646\text{-}674}$ | -30.6 ± 3.4 | -287.8 ± 24.4 | -4.7 ± 1.1 | 11 | -45 |
| $EIN3^{86\text{-}178}$-$EIN2^{646\text{-}674}$ | -28.6 ± 6.3 | -213.3 ± 6.3 | -1.1 ± 0.4 | 14 | -40 |
| $EIN3^{86\text{-}180}$-$EIN2^{646\text{-}674}$ | -35.1 ± 1.4 | -288.5 ± 44.4 | -1.6 ± 0.6 | 11 | -27 |
| $EIN3^{86\text{-}185}$-$EIN2^{646\text{-}674}$ | -25.2 ± 4.3 | -308.6 ± 75.5 | -5.5 ± 2.4 | 11 | -21 |
| $EIN3^{86\text{-}190}$-$EIN2^{646\text{-}674}$ | -30.4 ± 9.4 | -143.0 ± 22.3 | -9.3 ± 3.8 | 10 | -19 |
| $EIN3^{86\text{-}195}$-$EIN2^{646\text{-}674}$ | -19.7 ± 5.7 | -273.6 ± 41.8 | 15.0 ± 4.7 | 12 | -13 |
| $EIN3^{86\text{-}200}$-$EIN2^{646\text{-}674}$ | -22.4 ± 4.3 | -302.7 ± 81.7 | 14.1 ± 4.3 | 15 | -8 |
| $EIN3^{86\text{-}205}$-$EIN2^{646\text{-}674}$ | -39.5 ± 6.7 | 309.9 ± 5.4 | 26.0 ± 0.5 | 6 | 3 |
| $EIN3^{86\text{-}210}$-$EIN2^{646\text{-}674}$ | -30.1 ± 6.1 | -250.2 ± 17.6 | 15.0 ± 0.4 | 11 | 5 |
| $EIN3^{86\text{-}215}$-$EIN2^{646\text{-}674}$ | -46.3 ± 7.9 | -300.2 ± 7.2 | 28.0 ± 2.0 | 12 | 10 |

**Table S4. Information of the EIN2 and EIN3 sequences used in the multiple sequence alignment.**

| Protein | Species | Uniprot ID |
|---|---|---|
| EIN2<br>EIN3 | *Arabidopsis thaliana* | Q9S814<br>O24606 |
| EIN2<br>EIN3 | *Vitis vinifera* | F6HR32<br>A0A438JCI9 |
| EIN2<br>EIN3 | *Helianthus annuus* | A0A251UC58<br>A0A251SIP3 |
| EIN2<br>EIN3 | *Nicotiana attentuata* | A0A142IX47<br>A0A1J6HTH5 |
| EIN2<br>EIN3 | *Solanum tuberosum* | M1BY00<br>M1A5I7 |
| EIN2<br>EIN3 | *Solanum lycopersicum* | A0A3Q7HW43<br>Q94FV4 |
| EIN2<br>EIN3 | *Coffea arabica* | A0A6P6SYT8<br>A0A6P6TVG4 |
| EIN2<br>EIN3 | *Actinidia chinensis* | A0A2R6PWM2<br>A0A075EAR5 |
| EIN2<br>EIN3 | *Carica papaya* | AIO12155.1<br>A0A0R5PE43 |
| EIN2<br>EIN3 | *Musa acuminata* | 065046552.1<br>A1IIY3 |
